# Impacts of maternal microbiota and microbial metabolites on fetal intestine, brain and placenta

**DOI:** 10.1101/2022.07.01.498433

**Authors:** Aleksi Husso, Tiina Pessa-Morikawa, Ville Mikael Koistinen, Olli Kärkkäinen, Leo Lahti, Antti Iivanainen, Kati Hanhineva, Mikael Niku

## Abstract

The maternal microbiota modulates fetal development, but the mechanisms of these earliest host-microbe interactions are unclear. We compared full-term fetuses from germ-free (GF) and normally colonized mouse dams by gene expression profiling and non-targeted metabolomics. The developing immune system was strongly dependent on the maternal microbial status. In the fetal intestine, critical components mediating host-microbe interactions were differentially expressed. In fetal brain and placenta, interferon and inflammatory signaling were downregulated in germ-free fetuses. Neural system development and function, translation and RNA metabolism, and regulation of energy metabolism were significantly affected at the gene expression level. These impacts were strongly associated with microbial metabolite concentrations in the fetal tissues, suggesting that they are largely, although perhaps not exclusively mediated by maternal microbial metabolites absorbed through placenta. Several aryl sulfates were among the compounds strongly associated with gene expression differences. The germ-free fetus may suffer from depletion of queuine, a bacterial hypermodified nucleobase essential for eukaryotic tRNA stability and function.

## Introduction

The development and programming of the immune system, metabolism and the central nervous system requires interactions with the commensal microbiota (Ganal-Vonarburg et al., 2020). We are connected with our prokaryotic companions already before birth. The maternal microbiota stimulates the fetal generation of intestinal lymphoid cells, and the early host-microbe interactions induce tolerance to avoid excessive reactivity and inflammatory pathologies (Al Nabhani and Eberl, 2020; Gars et al., 2021; Gomez de Aguero et al., 2016). Absence of maternal microbiota during pregnancy predisposes the offspring to metabolic syndrome and affects the differentiation of enteroendocrine cells and the sympathetic nerves (Kimura et al., 2020). Maternal microbes also promote axonogenesis in the fetal brain and modulate the differentiation of microglia (Gars et al., 2021; Thion et al., 2018; Vuong et al., 2020).

Live bacteria are rare in a healthy fetus (Hornef and Penders, 2017; Husso et al., 2021). Thus, the prenatal host-microbe interactions are likely primarily mediated by circulating metabolites and other components of microbes, reaching the fetus through placenta. Metabolites generated or modified by gut microbiota penetrate all host tissues (Uchimura et al., 2018). We recently showed that thousands of microbially modulated metabolites are found also in the fetus, by non-targeted metabolomics comparison of fetuses from germ-free and conventional mouse dams (Pessa-Morikawa et al., 2022). A hundred compounds were undetectable in germ-free fetuses, indicating that their synthesis is dependent on the microbiota. We identified several metabolites with reported effects on host physiology and fetal development, such as 3-indolepropionic acid, trimethylamine N-oxide (TMAO) and 5-aminovaleric acid betaine (5-AVAB). Well-studied bacterially derived metabolites include the short chain fatty acids (SCFAs), produced by microbial metabolism of dietary complex carbohydrates, and microbially modified secondary bile acids (Li et al., 2022). SCFAs contribute to the fetal programming of energy metabolism and sympathetic nervous system development (Kimura et al., 2020). Experiments with monocolonized dams revealed the importance of microbial aryl hydrocarbons on the fetal immune system (Gomez de Aguero et al., 2016). The majority of microbial metabolites and especially their effects on mammalian fetal development are however still unknown (Ganal-Vonarburg et al., 2020).

Microbial metabolites are sensed by multiple receptor systems, including the SCFA-activated G-protein coupled receptors (GPCRs) and nuclear receptors such as aryl hydrocarbon receptor (AhR), pregnane X receptor (PXR), bile acid receptor alias farnesoid X activated receptor (FXR), and peroxisome proliferator-activated receptors (PPAR) (Ganal-Vonarburg et al., 2020; Klepsch et al., 2019; Kou and Dai, 2021; de Vos et al., 2022). These signaling pathways modulate immunity, energy metabolism and neurophysiology, and are critical in host health and development.

In this study, we investigate the impact of maternal microbial metabolites on fetal intestine, brain and placenta. We analyze gene expression profiles in these organs from germ-free and specific pathogen free mouse dams, and associate the gene expression data with our recent non-targeted metabolomics data (Pessa-Morikawa et al., 2022). Our observations indicate major impacts of maternal microbiota on fetal development, strongly associated with microbially modulated metabolites.

## Materials and methods

### Animals

Fetal and placental mouse organ samples from pregnant germ-free (GF, n=6) and specific pathogen free (SPF, n=6) C57BL/6J dams were obtained from the EMMA Axenic Service at Instituto Gulbenkian de Ciência, Portugal, as described in detail previously (Pessa-Morikawa et al., 2022). The dams were euthanized 18.5 days post coitum. Brain, intestine and placenta were collected from 4 fetuses per dam (two for gene expression profiling and two for metabolomics), for a total of 12 fetuses per experimental group per method. The fetal organ samples were frozen in liquid nitrogen after collection, stored at −80°C and shipped on dry ice. Both groups had 7 male fetuses and 5 female fetuses. The GF and SPF status of the dams were regularly monitored by culture and 16S qPCR. The GF dams were 3-4 months old and the SPF dams 4-5 months old. All dams were fed identical RM3-A-P breeding diets (SDS Special Diet Services, Essex, UK), autoclaved at 121 °C. The SPF feed was autoclaved for 20 minutes and the GF feed for 30 minutes due to logistical reasons.

### RNA extraction and gene expression profiling

Total RNA was extracted from fetal intestine, brain and placenta using the Qiagen RNeasy Mini Kit. The tissues were mechanically lysed by grinding with a plastic pestle attached to an electric drill, and the protocol included the optional on-column DNAse digestion.

Quality and quantity of the extracted RNA samples were analyzed with LabChip GX Touch HT RNA Assay Reagent Kit (PerkinElmer, Waltham, MA, USA) and Qubit RNA BR kit (Thermo Fisher Scientific, Waltham, MA, USA). For genomic DNA contamination measurement, Qubit DNA BR kit (Thermo Fisher Scientific, Waltham, MA, USA) was used. The intestinal RNA extracts were re-purified using the Qiagen RNeasy Micro Plus kit to remove the remaining genomic DNA; the brain and placental extracts did not require additional purification.

Dual-indexed mRNA libraries were prepared from 150 ng of total RNA with QuantSeq 3’ mRNA-Seq Library Prep Kit FWD (Lexogen Gmbh, Vienna, Austria) according to user guide version 015UG009V0251. During second strand synthesis, 6 bp Unique Molecular Identifiers (UMI) were introduced with UMI Second Strand Synthesis Module (Lexogen Gmbh, Vienna, Austria) for detection and removal of PCR duplicates. Quality of the libraries was measured with LabChip GX Touch HT DNA High Sensitivity Reagent Kit (PerkinElmer, Waltham, MA, USA). Sequencing was performed with NovaSeq 6000 System (Illumina, San Diego, CA, USA) with read length of 2×101 bp and target coverage of 10 M reads for each library.

QuantSeq 3’ mRNA-Seq Integrated Data Analysis Pipeline version 2.3.1 FWD UMI (Lexogen Gmbh, Vienna, Austria) on Bluebee® Genomics Platform was used for primary quality evaluation of the RNA sequencing data. The pipeline utilizes UMI sequences to remove PCR duplicates, STAR Aligner (Dobin et al., 2013) with modified ENCODE settings for alignment, and HTSeq-count (Putri et al., 2022) with QuantSeq FWD-specific options for gene read counting.

Differential gene expression (DE) analysis was performed using the QuantSeq FWD-UMI Data Analysis Pipeline, which utilizes DESeq2 (Love et al., 2014), for each tissue type, comparing between the SPF and GF groups.

### Metabolomics

The samples were analyzed by a liquid chromatography–mass spectrometry (LC-MS) system, consisting of a 1290 Infinity Binary UPLC coupled with a 6540 UHD Accurate-Mass Q-TOF (Agilent Technologies Inc., Santa Clara, CA, USA), as described previously (Pessa-Morikawa et al., 2022). Briefly, a Zorbax Eclipse XDB-C18 column (2.1 × 100 mm, 1.8 μm; Agilent Technologies) was used for the reversed-phase (RP) separation and an Acquity UPLC BEH amide column (Waters Corporation, Milford, MA, USA) for the HILIC separation.

The peak detection and alignment were performed as previously reported in Pessa-Morikawa et al., 2022 and described in Klåvus et al., 2020 by Afekta Technologies Ltd. (Kuopio, Finland). The metabolite annotations, focusing on molecular features missing from the GF mice (and thus likely representing microbial metabolites), were performed in MS-DIAL v4.70 based on in-house and publicly available spectral databases and in MS-FINDER v3.50 using *in silico* molecular formula and MS/MS fragmentation prediction. Putative annotations were obtained for 17 previously unannotated molecular features which were missing from GF fetuses (Pessa-Morikawa et al., 2022).

### Statistical analysis

Multiple hypothesis adjusted p-values for DE analyses were calculated using Benjamini Hochberg correction.

Volcano plots were generated using the R package *EnhancedVolcano* (Blighe et al., 2022).

Over-representation analyses (ORA) were performed using g:profiler version e105_eg52_p16_e84549f (Raudvere et al., 2019) and Metascape 3.5 (Dec 18, 2021 release) (Zhou et al., 2019), using gene priorization by evidence counting. Transcription factor binding sites were analyzed using Transfac (Matys et al., 2006) through g:profiler.

Gene set enrichment analysis (GSEA) by functional class scoring (Subramanian et al., 2005) was performed with the GSEA version 4.2.2 using Hallmark gene sets (50 gene sets), canonical pathways (2892 gene sets) and immunosignature gene sets (5219 gene sets) from Molecular Signatures Database (version 7.5.1). The expression datasets contained data of 18196 genes after collapsing to features to gene symbols. Limits for the number of genes in queries were set at 15 min and 500 max and phenotype permutations at 1000. Enriched gene sets with FDR < 25% were considered significant, following the recommendation of the Broad Institute for exploratory studies.

Associations between metabolomics and transcriptomics data (as averages of the two fetuses per dam) were explored in R version 4.1.2, using *phyloseq* version 1.38 (McMurdie and Holmes, 2013) and *microbiome* version 1.17.2 (Lahti and Sudarshan, 2022). Associations of individual molecular features and genes were evaluated by Spearman correlations, selecting pairs which showed ρ>0.9 in SPF mouse data for further analyses.

Hierarchical clustering was performed using the R core method *hclust (*complete linkage), using the dissimilarity matrix (1-Spearman). Associations between hierarchical gene and molecular feature clusters were evaluated by mean Spearman correlations of cluster members, and clusters with mean ρ>0.7 in the SPF group (for brain, ρ>0.6, due to the much smaller number of DEGs) were selected for further analyses. These contained, as averages per tissue, 16-43 genes or 7-15 molecular features per cluster.

Plaid Model biclustering was performed using *biclust* version 2.0.3 (Kaiser et al., 2022). The biclusters were generated with fit.model = y∼m from correlation matrix [abs(Spearman)] using the SPF mouse data only. Approximately ten biclusters were generated for each tissue. Biclusters with strongest mean internal correlations between molecular features and genes were selected for further analyses.

Molecular features were scored based on the order of numbers of significant hits in g:profiler over-representation analyses for their associated gene sets (combined score based on direct correlations, hierarchical cluster correlations, and biclustering).

The top-scoring molecular features in each tissue were selected for over-representation analyses of the strongly associated genes (Spearman ρ>0.9 in SPF mouse data). Heatmaps were generated using Metascape 3.5 with gene priorization by evidence counting, selective GO clusters, all available murine terminologies, and protein-protein interaction enrichment.

## Results

### Gene expression profiling

Differential expression (DE) analysis of germ-free (GF) and specific pathogen free (SPF) mice indicated a major impact of maternal microbiota on gene expression in fetal murine intestine (Table 1, Figs. 1 & 2 and Supplementary Table 1). A higher number of genes was downregulated than upregulated in GF fetal intestine (Fig. 2). Of the genes with Gene Ontology (GO) annotations for immune system, 75% were downregulated in the GF intestine; in contrast, 80% of the genes annotated for translation, ribosomes and tRNA metabolism were upregulated. The most significantly downregulated genes included several critical components in intestinal host-microbe interactions and immunity (*Pigr, Muc3, Sectm1b, Trim12c, Ace)*, transporters of the solute carrier family (*Slc39a4* and *Slc13a2*) and *Igfals*, a component of the growth control system. Multiple interferon and cytokine signaling genes, several complement genes and MHC-I genes, nine other antiviral *Trim* genes, antimicrobial lectins (*Reg3b, Reg4, Lgals3, Lgals9*) and the antimicrobial peptide *Ang* were also significantly downregulated in the GF intestine, along with the enteroendocrine cell markers *Insl5, Pyy, Gip* and *Nts*. The genes upregulated in GF fetal intestine included those coding for metallothionein-1, ribosomal and mitochondrial proteins, and the imprinted maternally expressed developmental noncoding RNA H19.

**Table 1.** Summary of differential expression analysis of GF vs SPF mouse fetal tissues.

| Tissue | Total genes represented<br>(>0 total reads) | Downregulated in GF<br>(p.adj < 0.1) | Upregulated in GF<br>(p.adj < 0.1) |
| --- | --- | --- | --- |
| Fetal intestine | 25599 | 1642 (6.4%) | 1352 (5.3%) |
| Fetal brain | 26839 | 204 (0.76%) | 185 (0.69%) |
| Placenta | 26282 | 547 (2.1%) | 470 (1.8%) |

**Figure 1.**
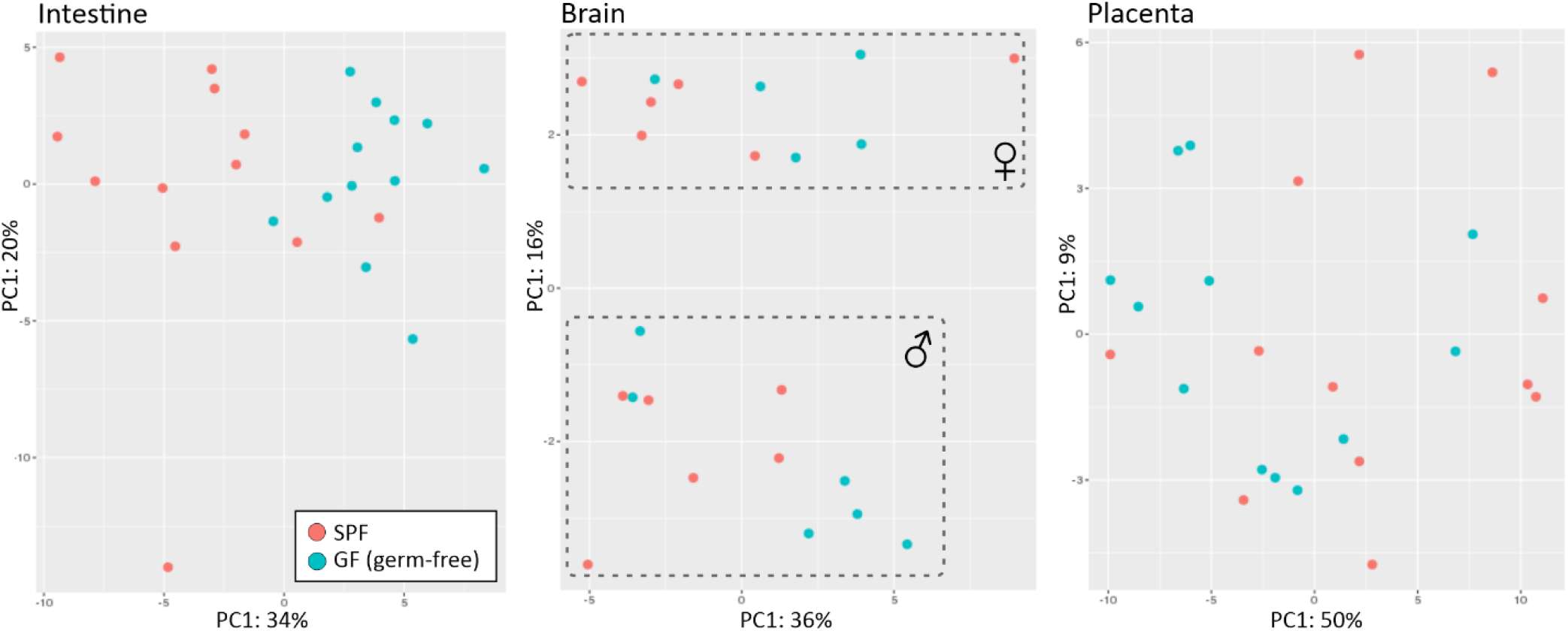
Principal component analysis (PCA) of gene expression profiles in fetal murine intestine, brain and placenta. The first two components of the normalized counts of expressed genes are shown. In fetal brain, clustering on sex was observed.

**Figure 2.**
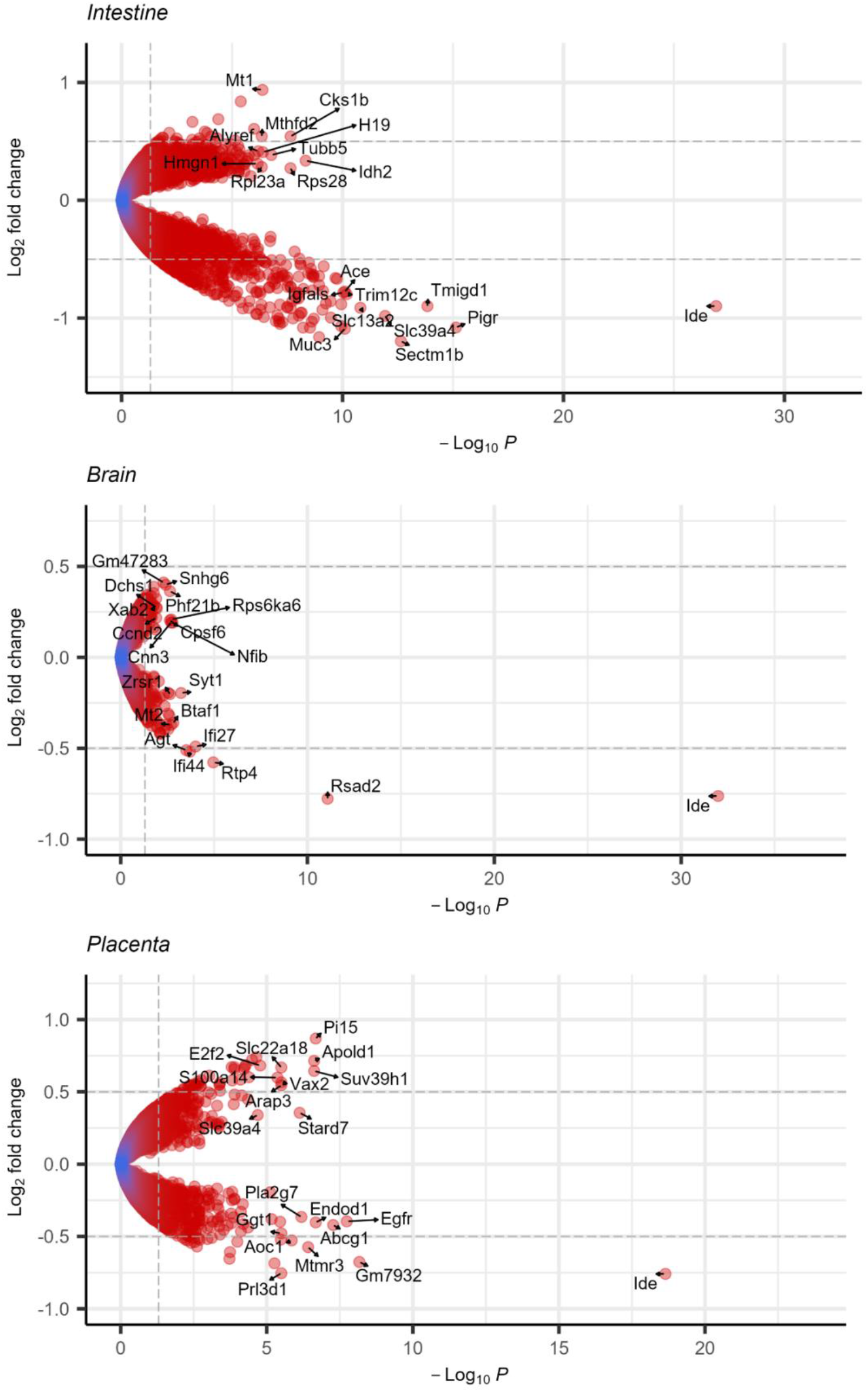
Volcano plots of differential gene expression in fetal intestine, fetal brain and placenta. Genes with negative Log_2_ fold change values were downregulated in GF mice vs SPF mice. Ten most significantly downregulated and upregulated genes are labelled. Dashed lines indicate Log_2_ fold change 0.5 and - Log_10_ adjusted P values.

In fetal brain and in placenta, the differences in gene expression profiles were clearly less prominent. Also there, a majority (71% and 58%) of immunity genes were downregulated in GF fetuses; genes for translation or neural system were quite evenly down- or upregulated. In PCA of fetal brain expression profiles, the fetuses actually clustered primarily by sex, which was not observed in intestine or placenta. All genes with significantly different expression levels in female and male fetal brain were X or Y chromosomal and included one Y chromosomal lncRNA (in the order of decreasing significance, these were: *Ddx3y, Kdm5d, Uty, Eif2s3y, Xist, Gm29650, Eif2s3x, Kdm6a, Kdm5c, Ddx3x* and *Pbdc1*). Eight out of ten statistically most significant DEGs were downregulated in GF fetal brain, including multiple interferon-related genes (*Rsad2, Ifi27, Ifi44, Rtp4;* Figure 2); significantly downregulated neural immunity genes also included *CX3CL1*. Interestingly, a component of the angiotensin system (*Agt*) was significantly downregulated also in GF brain. Multiple genes involved in neuronal development and synaptic signaling were significantly differentially expressed. Several genes involved in synaptic signaling (two synaptotagmins, *Shisa6*, two calmodulins and *Nrgn*) and the glia-specific *Gfap* and *Olig2* were significantly downregulated in the GF fetal brain. Genes upregulated in GF brain included those coding for neural stem cell regulator Phf12b, the neural transcription factors and activators Nfib, Sox11 and Eomes, and several cadherins involved in neural system development. Two spliceosome components (*Xab2* and the paternally expressed *Zrsr1*) were downregulated in GF fetuses. One brain sample was excluded from analyses as an outlier, as expression profiling identified it as non-brain head tissue.

Also in placenta, seven out of ten statistically most significant DEGs were downregulated in GF mice (Fig. 2). These included *Egfr*, the exosomal *Endod1*, the prolactin precursor *Prl3d1*, and the *Gm7932* gene coding for a lncRNA. The methyltransferase *Suv39h1*, the endothelial signaling protein *Apold1*, the trypsin inhibitor *Pi15* and two solute carriers were among the most significantly upregulated genes in the GF placenta.

Five genes were observed to be significantly differentially expressed in all three tissues: *Ide, Btaf1, Rhou, Rtp4* and *Tub4a*. All these were upregulated in SPF fetuses vs GF fetuses, except for *Tub4a* in brain.

### Functional enrichment analyses

GSEA was used to identify Hallmark gene sets enriched among the differentially expressed genes in fetal brain, intestine and placenta samples. The interferon alpha and interferon gamma response gene sets were the most positively enriched in SPF fetal intestine and brain as compared to GF, with -Log10 q >1.5 (Fig 3. A and B) corresponding to FDR < 5%. Heat maps for these gene sets for fetal intestine and brain are shown in Supplementary Fig. 1. The leading-edge genes in these gene sets included *Irf7, Irf9* and *Stat2* transcription factors, components of the MHC complex, *Ifi44* and *B2m*, and interferon stimulated genes (ISGs) such as *Rsad2, Trim5, Oas1, Oasl*. In placenta, the most positively enriched Hallmark gene set in SPF mice was IL6-JAK-STAT-signaling with -Log10 q 1.22 corresponding to FDR 6% (Fig.3 C). In all, 20 positively enriched Hallmark gene sets were significant at FDR < 25% in fetal intestine, including bile acid metabolism, estrogen response early/late, inflammatory response and various cytokine signaling pathways (Fig. 3 A). 18 positively enriched Hallmark gene sets were significant in fetal brain and three in placenta at FDR <25%.

**Figure 3.**
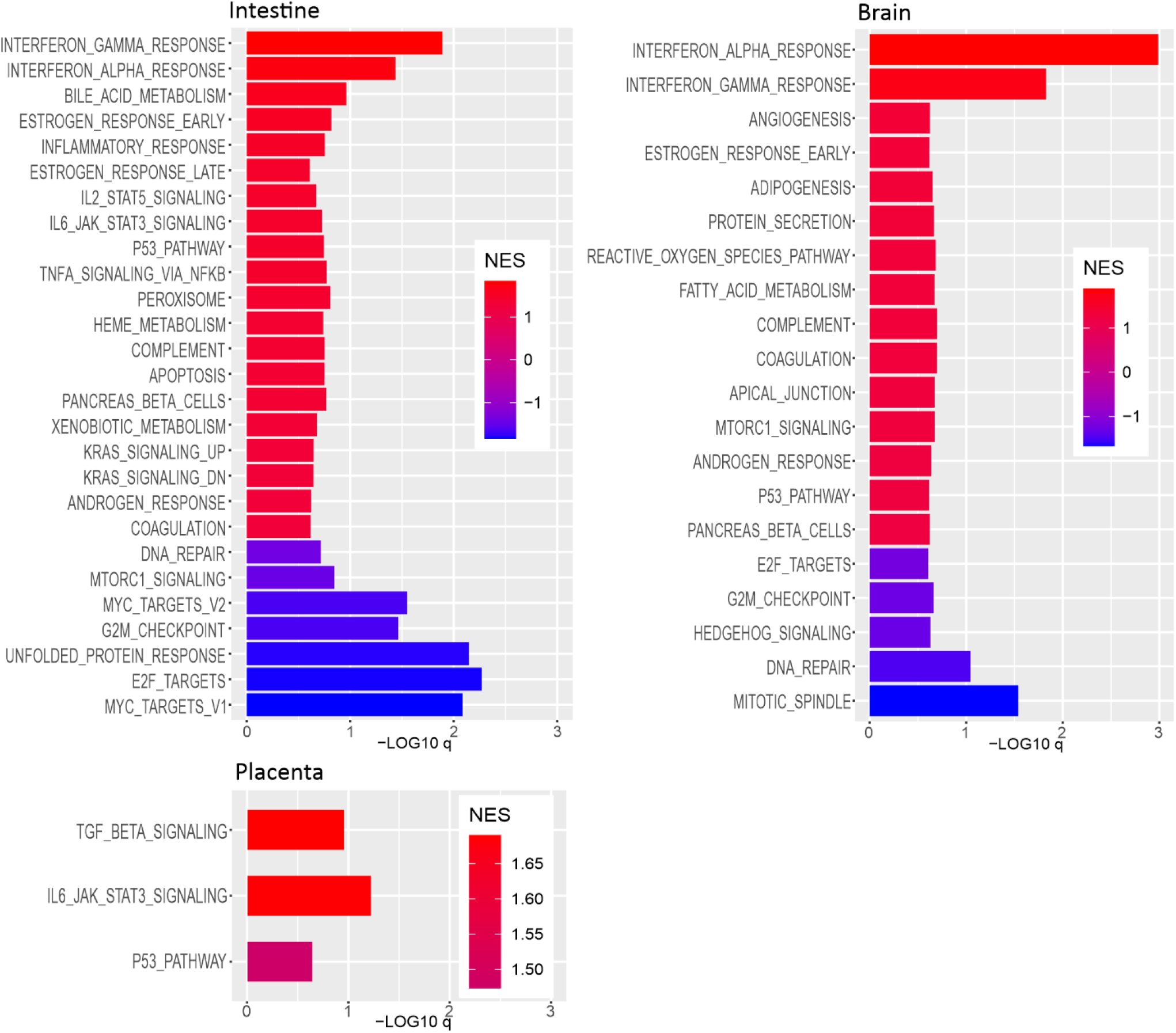
Hallmark gene sets enriched at FDR <25% in gene set enrichment assay in fetal intestine, brain and placenta. -Log10 q of gene sets with positive (red bars) and negative (blue bars) net enrichment score (NES) for specific pathogen free (SPF) versus germ-free (GF) fetuses.

7 Hallmark gene sets in fetal intestine and 5 sets in fetal brain were significantly negatively enriched at FDR < 25% in SPF versus GF mice (Fig 3. A-B, blue bars). These included proliferation-associated gene sets, such as Myc targets, E2F targets and mitotic spindle. No significantly enriched Hallmark gene sets in GF over SPF mice were detected in placenta.

Enrichment of gene sets related to interferon signaling and virus response was also seen in SPF vs GF fetal intestine and brain, but not in placenta, by GSEA analysis using the C2 canonical pathways and C7 immunosignature gene sets (Supplementary table 2).

Over-representation analysis (ORA) of DEGs (p.adj < 0.05) suggested strong impact of maternal microbiota on the fetal immune system in all tissues analysed (Fig. 4; for more details, see Supplementary Figure 2 and Supplementary Table 3). Especially in fetal brain, this was most marked in pathways related to interferons and virus responses.

**Figure 4.**
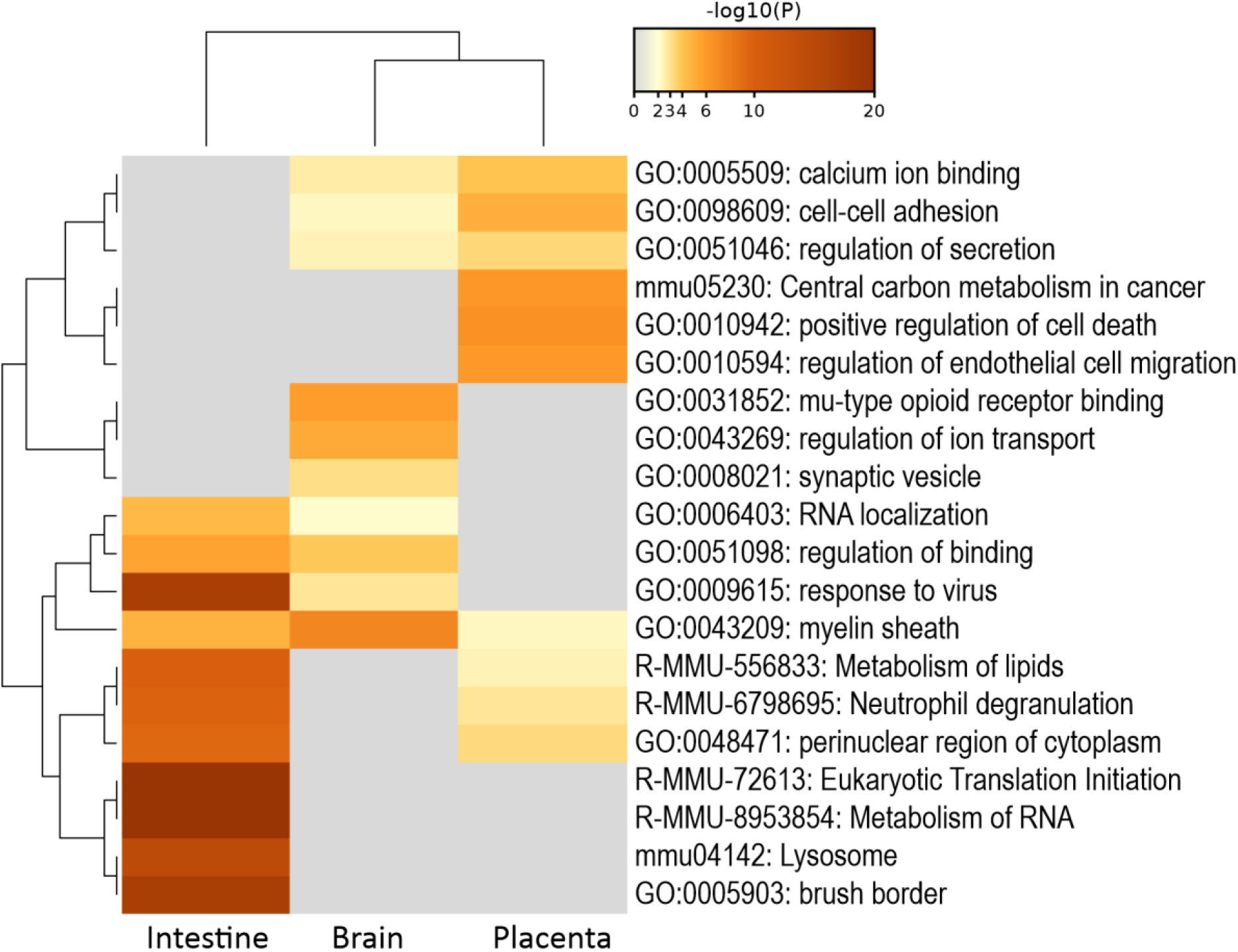
Over-representation analysis of genes which were significantly differentially expressed (q < 0.05) in GF versus SPF fetal murine intestine, brain and placenta. Top 20 enriched ontology terms are shown; for more details, see Supplementary Figure 2.

In the fetal intestine, genes related to translation, RNA and lipid metabolism, and brush border / absorption were also significantly over-represented among DEGs. Genes downregulated in GF intestine were enriched for immune system, lipid metabolism and membrane transport, including genes coding for the multidrug resistance protein Abcc2 (Mrp2) and the ABC sterol transporters Abcg8 and Abcg5. Genes upregulated in GF fetuses were enriched for translation, RNA metabolism and cell cycle/mitosis, and included several subunits of the translation initiation factor eIF2 and the elongator acetyltransferase complex.

In fetal brain, ORA indicated significant differences in neuronal synapse and opioid receptor pathways. Genes downregulated in GF fetuses were enriched for interferon responses, synaptic processes and regulation of ion transport; CDR-mediated mRNA stabilization was most significantly over-represented among genes upregulated in GF animals.

In placenta, pathways in development and morphogenesis (especially of blood vessels), proteolysis, and cell death were primarily over-represented. Genes downregulated in GF placenta were most significantly enriched for positive regulation of cell death and tube morphogenesis. Upregulated genes were enriched for myeloid homeostasis and transport of small molecules.

Regarding predicted transcription factor binding sites, the DEGs in the three tissues were most significantly enriched for E2Fs, Sp-1 and Usf. Predicted targets of interferon regulatory factors (IRFs) and early growth response proteins (EGRs) were also significantly enriched. DEGs in fetal intestine (but not in brain and placenta) were significantly enriched for multiple predicted binding sites of aryl hydrocarbon receptor (AhR) and AhR nuclear translocator (Arnt). DEGs in intestine and placenta (but not in brain) were also significantly enriched for some binding sites for the vitamin D receptor (VDR) and/or the farnesoid X activated receptor / bile acid receptor (FXR).

### Associations between gene expression profiles and metabolomics data

To explore potential effects of microbial metabolites on fetal intestine, brain and placenta, we analysed associations between metabolites and gene expression, focusing on metabolites which were significantly more abundant in SPF fetuses compared to GF fetuses, or undetectable in GF fetuses. To detect various types of associations between individual compounds and genes, and between molecular feature groups and gene families or co-regulated pathways, we evaluated 1) correlations of individual metabolites with individual genes, 2) correlations of clusters of metabolites with clusters of genes (by hierarchical clustering), and 3) correlations within biclusters composed of metabolites and genes. In addition to a significant difference between the experimental groups, we required strong correlation within the SPF group (Spearman ρ>0.9), in order to focus on metabolite-gene-associations which are likely due to actual abundances of the compounds, rather than due to the overall differences of SPF/GF physiology.

All molecular features were scored based on significant hits in over-representation analyses of the strongly associated genes or gene clusters (Supplementary Table 4). Highest-scoring molecular features were then examined in more detail by over-representation analyses of genes strongly associated with them. These associations are reported here (Figs. 5-7; for more detail, see Supplementary Figures 3-5).

**Figure 5.**
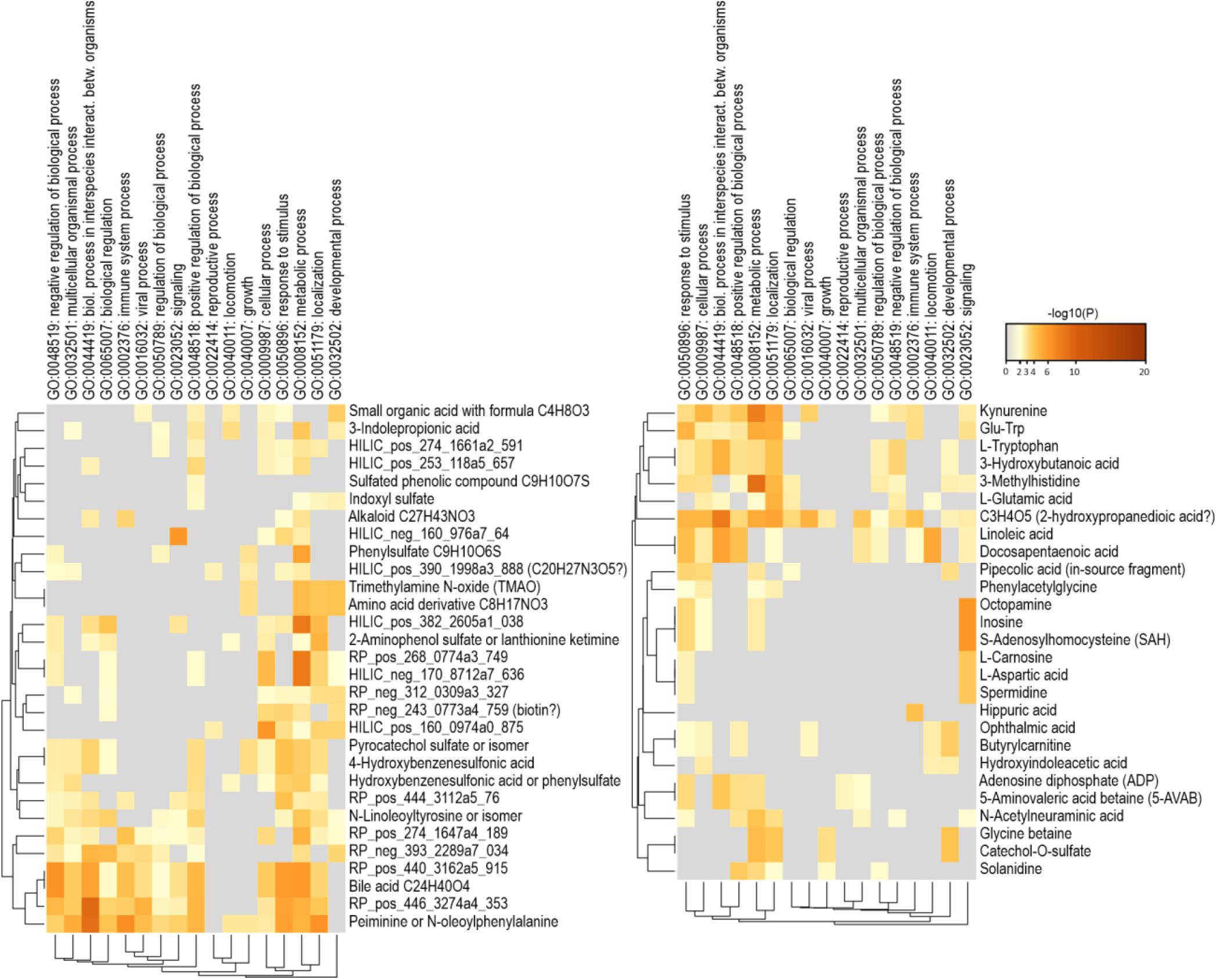
Over-representation analysis of genes strongly associated with metabolites in fetal intestine. Highest-scoring molecular features missing from GF fetuses (left) and highest-scoring annotated metabolites more abundant in SPF fetuses (right).

**Figure 6.**
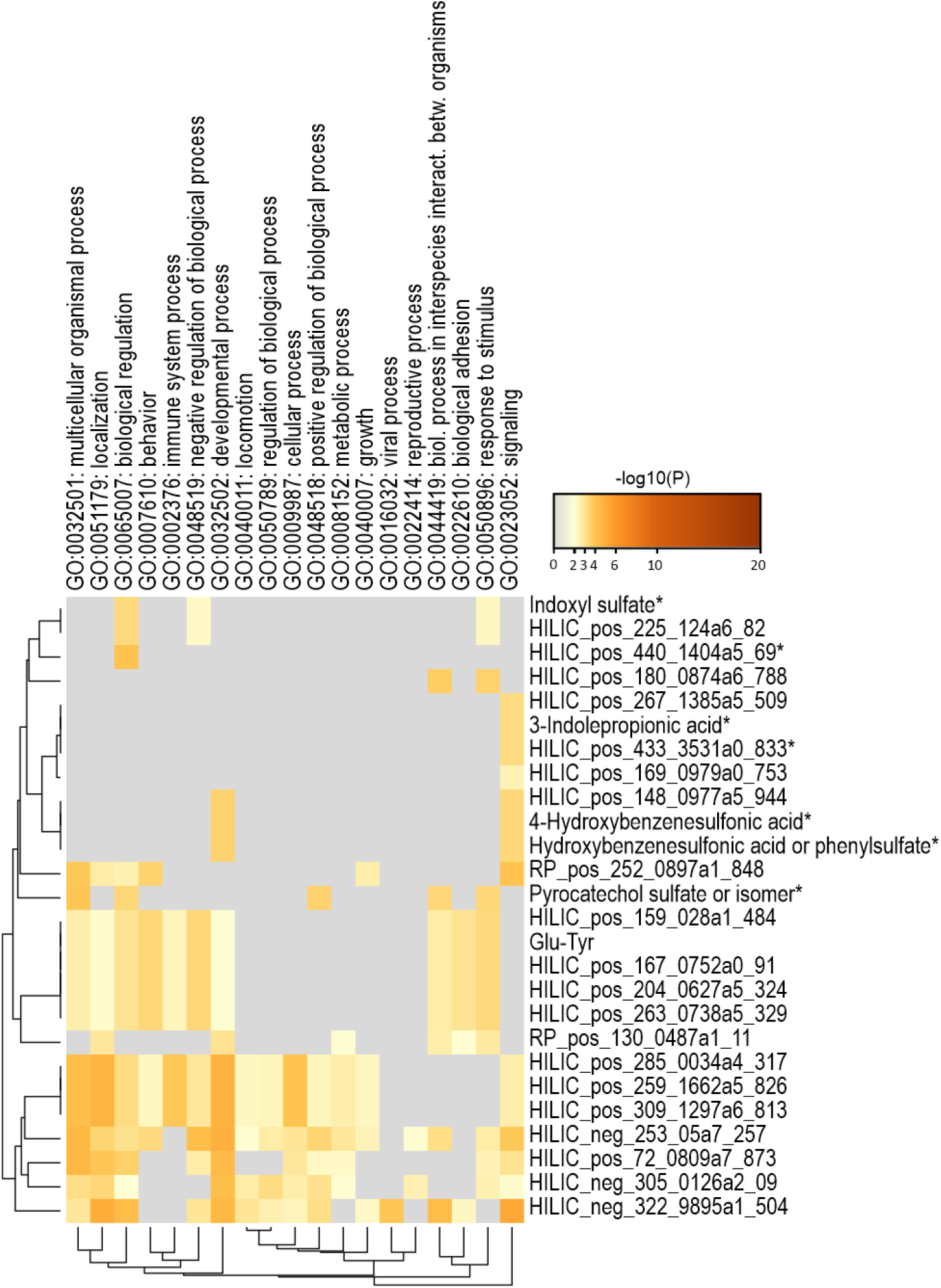
Over-representation analysis of genes strongly associated with metabolites in fetal brain. Metabolites missing from GF fetuses are indicated by *.

**Figure 7.**
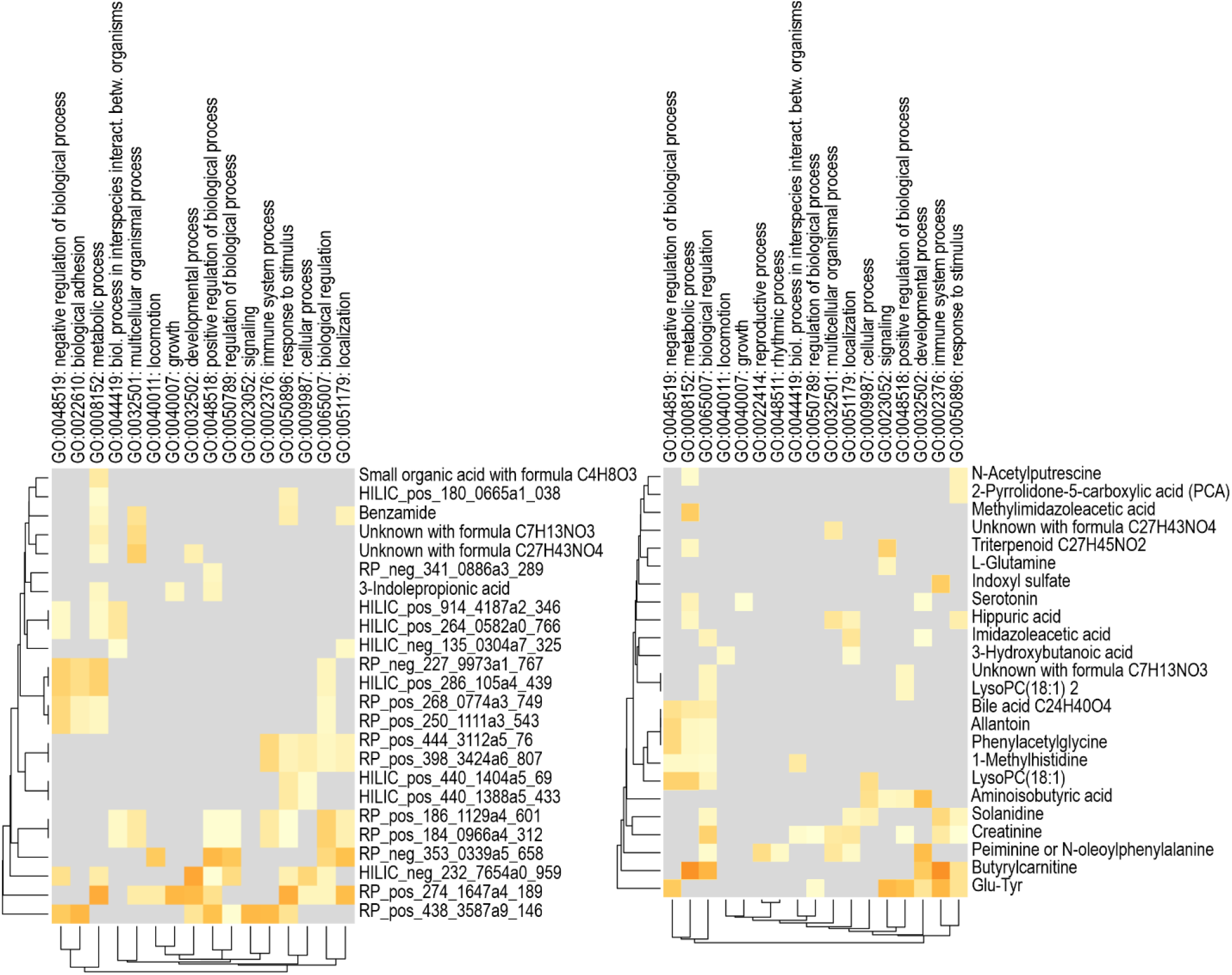
Over-representation analysis of genes strongly associated with metabolites in placenta. Highest-scoring molecular features missing from GF fetuses (left) and highest-scoring annotated metabolites more abundant in SPF fetuses (right).

Examples of the strongest associations between individual molecular features and genes are shown as scatterplots in Fig. 8.

**Figure 8.**
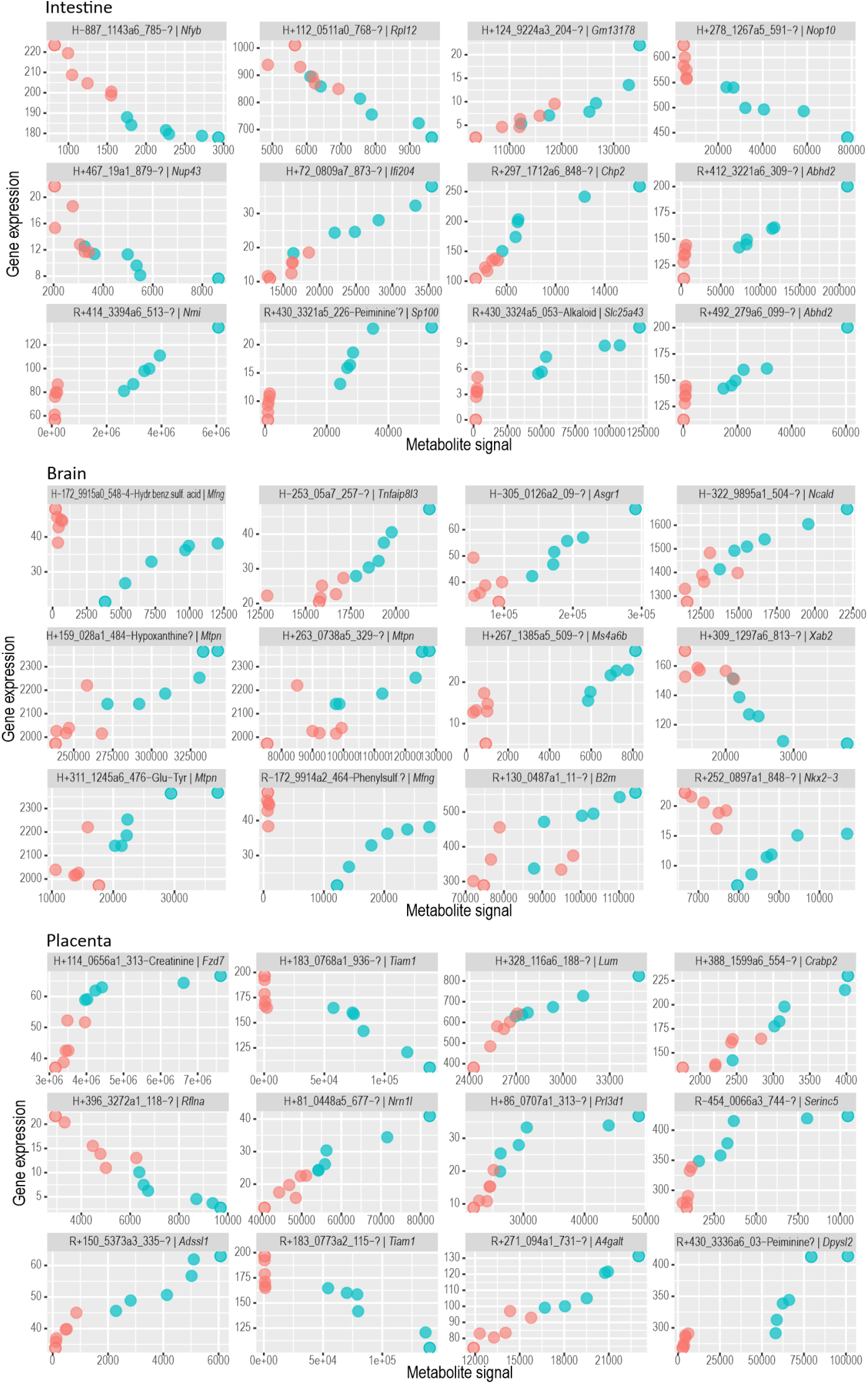
Strongest associations between metabolites and gene expression in fetal murine intestine, brain and placenta. Only one gene per metabolite is shown. Molecular feature signal intensity on X axis, normalized gene expression on Y axis. Each point is the mean of the fetuses from one dam. Red = germ-free, blue = specific pathogen free. Unidentified or tentatively annotated metabolites are marked with “?”. R = RP, H = HILIC.

In fetal intestine and placenta, >95% of significantly differentially expressed genes (p.adj < 0.05) were strongly associated with at least 3 molecular features (Spearman ρ>0.9 in the SPF group). In brain, 31% of DEGs showed such associations. More than 10 molecular feature associations were observed for 58%, 66% and 8% of DEGs in fetal intestine, placenta and brain, and more than 50 associations for 10%, 12% and 0%, respectively. In brain, genes with Gene Ontology annotations for immunity or neurophysiology had more metabolite associations (approximately 70% of these genes had at least 3 associations, and 20% had more than 10).

In intestine and placenta, expression levels of immunity genes (by Gene Ontology annotations) were mostly positively correlated with metabolomics signal intensities. In brain, such differences were not observed. In contrast, genes associated with translation correlated mostly negatively in the fetal intestine.

A total of 148 molecular features which were not detectable in GF fetuses showed associations with gene expression profiles (Supplementary Table 4). 84 of these were also missing from GF placentae. 23 of the molecular features missing from GF fetuses were at least putatively characterized (identification level 3 or better).

Six of these were aryl sulfates (4-hydroxybenzenesulfonic acid, indoxyl sulfate, pyrocatechol sulfate, and phenylsulfates). These were among the compounds with strongest associations to gene expression patterns in all tissues investigated. In the over-representation analysis of fetal intestine, these were associated with immunity (response to biotic stimulus and activation of innate immune response), lipid metabolism, regulation of cell growth, aromatic compound biosynthesis and tRNA metabolism. In brain, aryl sulfates associated with viral infection pathways, dopaminergic synapse, RAS signaling, and response to organocyclic compound. In placenta, they did not show significant associations.

3-indolepropionic acid and two unknown compounds (all observed only in SPF fetuses) were associated with adaptive immune system and Ras signal transduction in brain. In the fetal intestine, 3-indolepropionic acid was primarily associated with RNA metabolism, and in placenta, with regulation of cell growth.

Kynurenine, tryptophan and 3-methylhistidine were associated with immunity (largely positively) and RNA metabolism (mostly negatively) in the intestine. 1-methylhistidine was associated with virus response in placenta. Several other amino acids and their derivatives were also significantly more abundant in the SPF fetuses.

Tryptophan derivatives and aryl sulfates are typical AhR ligands. As fetal intestinal DEGs were enriched for predicted AhR/Arnt targets, we specifically looked at associations of such annotated compounds with transcription factor binding sites. Genes strongly associated with tryptophan, kynurenine, indolepropionic acid, a phenylsulfate and several compounds with tentative automatic annotation as tryptamine and indoleacetic acids were significantly enriched for predicted AhR/Arnt binding sites. This was not observed for indoxyl sulfate, hydroxyindoleacetic acid, 4-hydroxybenzenesulfonic acid and pyrocatechol sulfates. Instead, indoxyl sulfate, 4-hydroxybenzenesulfonic acid, pyrocatechol sulfates, and compounds automatically annotated as indoleacetic acids were associated with predicted VDR targets.

The betaine trimethylamine N-oxide (TMAO; only detected in the SPF fetuses) was associated with brush border, absorption and lipid metabolism in the SPF fetal intestine. In brain the only strong correlation (positive) was with the *Ide* gene. In placenta, there were no significant ORA hits. 5-aminovaleric acid betaine (5-AVAB) was associated with response to biotic stimulus and lipid metabolism in the intestine.

Butyrulcarnitine and several unannotated molecular features were associated with immunity (response to biotic stimulus, virus response, neutrophil degranulation, interleukin-1 production) and lipid metabolism in the intestine. Also in placenta, butyrulcarnitine was associated with neutrophil degranulation, and with oxidoreductase and peroxidase activities.

A bile acid (with retention time matching the secondary bile acid chenodeoxycholic acid), N-linoleyltyrosine or its isomer, and peiminine or N-oleoylphenylalanine (putative classifications) and two unannotated features were strongly associated with multiple pathways in the intestine. These included immunity (response to biotic stimulus, innate immunity, pattern recognition receptor signaling, T lymphocyte differentiation, several virus response pathways), translation, brush border and intestinal absorption. These compounds showed no significant ORA hits in brain or placenta.

An alkaloid (C27H43NO3) was associated with virus response and antigen processing & presentation in the intestine.

Two putatively classified features (2-aminophenol sulfate or lanthionine ketimine, and an amino acid derivative C8H17NO3) were associated with brush border, absorption and lipid metabolism in the intestine.

A small organic acid (possibly hydroxybutyric acid) was associated with neutrophil degranulation in the intestine.

The dipeptides Glu-Trp and Glu-Tyr were associated with extracellular response stimulus, virus responses and other immunity pathways in all tissues investigated.

S-adenosylhomocysteine (SAH), octopamine and inosine were associated with mitochondrial pathways and glucose starvation response in the intestine.

In placenta, creatinine was associated with multiple immune response pathways, and allantoin with external stimulus response, cell differentiation and adhesion.

The largest number of direct associations with genes (306 genes with Spearman ρ>0.9 in SPF mouse intestine data) were observed for an unidentified metabolite with the probable formula C3H4O5 (possibly 2-hydroxypropanedioic acid alias tartronic acid), which was not detected in GF intestine, but was present in brain and placenta in both experimental groups. In over-representation analysis for fetal intestine, the associated genes were significantly enriched for immunity pathways (response to virus, cytokine signaling and adaptive immune response).

Several unannotated molecular features were associated with virus response, translation and ribonucleoprotein pathways in the intestine. One of these (RP_pos_278_1254a1_34) matched the MS1 mass of queuine, was undetectable in GF intestine and placenta and significantly less abundant in GF brain, and was associated with genes primarily enriched for translation and its initiation, ribosomes (including the aminoacyl-tRNA binding protein Rpl8), protein-L-isoaspartate (D-aspartate) O-methyltransferase activity, nonsense-mediated decay and E2F-1 targets in the intestine. In brain, there were multiple unannotated molecular features which were more abundant in SPF fetuses and associated with neural development, synaptic function and interferon / virus responses. In placenta, several unidentified compounds were associated with immune responses (inflammatory response, cytokine production and phagocytosis) in placenta.

Some metabolites showed an unexpected correlation pattern in GF vs SPF mice: genes which correlated *positively* with these metabolites in SPF fetuses were *upregulated* in GF fetuses (although they were present at low quantity or undetectable in these animals), and genes which correlated *negatively* in SPF fetuses were *downregulated* in GF fetuses (two examples can be seen in Fig. 8, fetal brain). Thus, although immunity-related genes were typically downregulated in GF fetuses, they often correlated *negatively* with these metabolites in the SPF fetuses. This pattern was observed for all identified aryl sulfates in both fetal tissues; in the fetal intestine for bile acids, TMAO, alanine and glycine betaine (but not 5-AVAB), pipecolic, ophthalmic and hippuric acid; and in the brain (but not in intestine) for 3-indolepropionic acid and N-acetyl-L-ornithine. In placenta, this pattern was less clear but was also observed for some aryl sulfates, alanine betaine, bile acids, pipecolic and ophthalmic acid, butyrylcarnitine, inosine, serotonin, 2-pyrrolidone-5-carboxylic acid (PCA) and solanidine. Such patterns were more common in brain (53% of all strong correlations) than in the intestine (17%) or in placenta (32%). They were more common with compounds which were undetectable in GF fetuses compared to those found in both animal groups (in brain, 74% vs. 44%; in intestine, 24% vs. 17%, and in placenta, 35% vs. 32%).

In the fetal intestine, genes with unexpected expression GF fetuses were enriched for negative regulation of innate immune response, negative regulation of Notch signaling, and lipid metabolism, whereas genes with expected expression were enriched for positive regulation of defense response, complement activation and translation (Fig. 9). This was not observed in brain or placenta.

**Figure 9.**
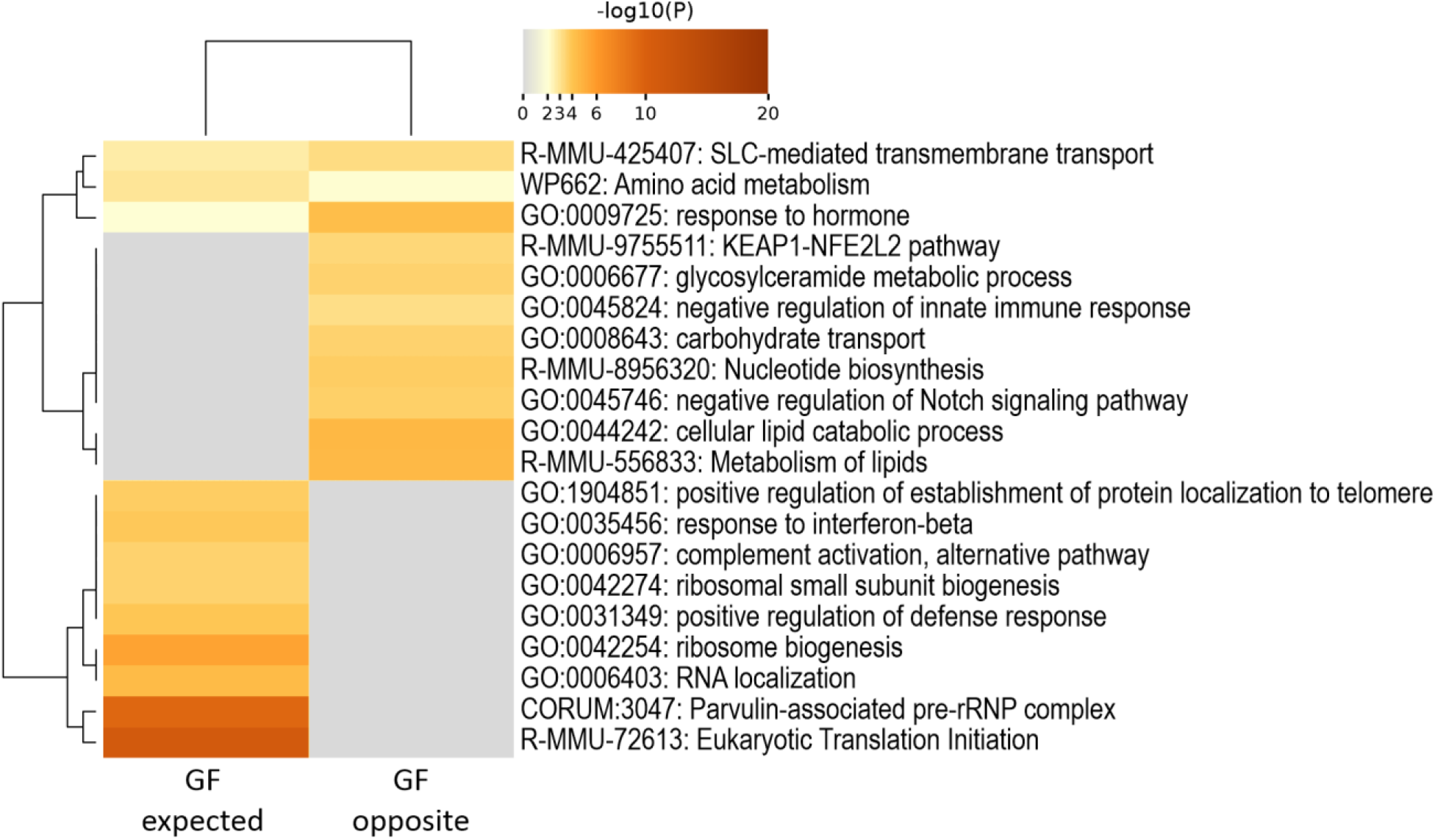
Heatmap of over-representation analysis of genes strongly associated with metabolites in fetal intestine. GF expected = expression in GF fetuses as expected from SPF fetuses; GF opposite = expression in GF fetuses different from that expected from SPF fetuses.

## Discussion

Our comparison of fetal intestine, brain and placenta from germ-free and specific pathogen free murine dams reveals major impacts of maternal microbiota on mammalian prenatal development, potentially largely mediated by circulating microbial metabolites. The maternal microbial status had a strong effect on the fetal immune system, translation, metabolism and neurophysiology. The effects were strongest in the fetal intestine. We observed multiple metabolites of likely microbial origin which were strongly associated with these processes in the fetus. Several of these have not been previously characterized in this context.

### Fetal intestine

In the fetal intestine, critical components for host-microbe interactions, innate immunity and epithelial barrier were affected by the maternal microbial status. Genes coding for mucins and the polymeric immunoglobulin receptor were among the most significantly downregulated in the GF fetal intestine, and expression of antimicrobial peptides and lectins was also suppressed. Inflammation, interferon responses and interleukin signalling were also downregulated in GF fetuses.

Secreted and membrane-bound mucins form the interface for host-microbe interactions in the intestine (Paone and Cani, 2020). The structure of the mucin layers is modulated by gut microbiota (Johansson et al., 2015; Szentkuti et al., 1990), but mucin expression begins early in fetal development (Buisine et al., 1998; Ferretti et al., 2016; Reid and Harris, 1998). Several mucin genes (and the associated trefoil family factors) were downregulated in our germ-free murine fetuses. Previous studies in adult mice have yielded conflicting results, with germ-free mice showing either downregulation (Larsson et al., 2012) or upregulation (Comelli et al., 2008) of mucins when compared to microbially colonized animals.

The polymeric immunoglobulin receptor (pIgR) transports IgA across the intestinal epithelium to the apical surface and is cleaved to generate the secretory component (Johansen and Kaetzel, 2011). In adult mice, pIgR expression is modulated by gut microbiota, possibly by TLR- and MyD88-mediated signalling. It is however expressed already early in fetal development in intestinal and other epithelia (Ben-Hur et al., 2004). We observed significant pIgR downregulation in the GF fetal intestine.

The antimicrobial peptide angiogenin (Sun et al., 2021) and C-type lectins Reg4 (Qi et al., 2020) and Reg3b (Bajic et al., 2020) regulate gut microbiota composition by selectively inhibiting e.g. α-proteobacteria, *Enterobacteriaceae*, or gram-negative bacteria, respectively. Reg3b is induced by the microbial metabolite propionate (Bajic et al., 2020).

The differentially expressed interferon response genes included IFN signal transduction mediators, notably *Irf7, Irf9* and *Stat2* that are key transcription factors in transcriptional activation of IFN downstream effectors (Lazear et al., 2019), components of the MHC complex, such as *Ifi44* and *B2m*, and numerous characterized interferon-stimulated genes (ISGs) such as *Rsad2, Trim5, Oas1* and *Oasl* that exhibit various types of anti-viral effector activities (Schoggins and Rice, 2011).

High representation of interferon-inducible genes among differentially expressed genes has been reported earlier in GF compared to conventional piglets (Sun et al., 2018). In their study, Sun et al. found seven genes belonging to this category out of total 14 DE genes which had lower expression in various tissues, including jejunum and colon, of GF animals compared to conventional controls. Four of these genes, *Ifit1, Rsad2, Ube2l6, Usp18* were among the genes with lower expression in fetal GF mouse intestine found in our study.

The downregulation of these virus response pathways in GF animals may be due to presumably lower exposure to viruses, or indirectly due to the lack of bacterial immune activation. Germ-free or antibiotic-treated mice have been shown to be more susceptible to many virus infections than conventional mice, and antibiotics induced a downregulation of interferon response genes in mouse pups (Garcia et al., 2021; Robinson and Pfeiffer, 2014). Gut microbes are known to modulate the host response to systemic viruses, such as the influenza virus (Winkler and Thackray, 2019). This may be due to tonic signals provided by the microbiota that set the homeostatic levels of the immune system components, including type 1 IFN, and calibrate the type 1 IFN response during viral infection. Potential mechanisms include activation of cells of the immune system through engagement of the pattern-recognition receptors (PRR) by binding of microbial molecules like lipopolysaccharides (LPS) (Winkler and Thackray, 2019)). Recent observations suggest a role for small-molecular microbial metabolites, such as secondary bile acids (Winkler et al., 2020).

Our observations complement and extend previous research on the prenatal effects of maternal microbiota on intestinal immunity and permeability (Gomez de Aguero et al., 2016). In that study, monocolonization of germ-free murine dams only during pregnancy significantly impacted the intestinal expression of *<u>Pigr</u>, Reg3b, Tmigd1, Slc13a2, H19* and 188 other genes common to our study in the offspring, analyzed two weeks after birth; for 76% of these, the change was in the same direction. In another study, antibiotic treatment of neonatal mice induced significant changes in the distal small intestinal expression of *Sectm1b, Slc39a4, Trim12c, Ace, Igfals* and 236 other genes common to our study, 93% of which in the same direction (Garcia et al., 2021). In contrast to the study by Gomez de Aguero et al., these also included multiple interferon signaling genes common to our data, all downregulated in the antibiotic treated pups.

Translation and RNA metabolism genes were upregulated in the GF fetal intestine. Gene expression profiling did not indicate obvious rationale for an increased protein synthesis, although mitosis and stress response pathways were over-represented among genes upregulated in the GF intestine (Zadjali, 2015). In adult GF intestine, proliferation is suppressed, and the characteristic enlargement of caecum is secondary to mucus and fiber accumulation. The upregulation of the components of the translation machinery may be a compensatory response to depletion of queuine, a hypermodified nucleobase exclusively synthesized by bacteria but essential to tRNA stability and function in eukaryotes (Fergus et al., 2015). Depletion of modified tRNAs is associated with pauses, misincorporation of amino acids and +1 frame shifts during translation, leading to protein misfolding and aggregation (Klassen et al., 2020). We detected a metabolite matching the MS1 mass of queuine, which was undetectable in GF intestine and placenta and significantly less abundant in GF brain, and associated with the expression of genes for translation, ribosomes, protein repair and nonsense-mediated decay in the intestine. Many of the genes for proteins affected by queuosine depletion in cell culture (Tuorto et al., 2018) and multiple genes linked to human tRNA modopathies (Chujo and Tomizawa, 2021) were significantly differentially expressed in the GF fetal intestine. These observations suggest that due to the lack of gut microbiota, the GF dam may be unable to provide sufficient queuine for the fetus, which then utilizes all available queuine (originating from dam diet) in queuosine synthesis.

### Fetal brain

In the fetal brain, maternal microbial effects were much more limited than in the intestine, likely due to the blood-brain barrier restricting the entry of microbial metabolites (Zhao et al., 2022). In fact, the fetal brain gene expression profiles clustered more clearly by sex rather than maternal microbial status. This was due to strong difference in the expression levels of several X and Y chromosomal genes, and the less prominent difference between GF and SPF fetuses, in comparison to the other tissues. However, gene expression analysis did indicate a substantial impact on brain immunity and neurodevelopment.

Many genes belonging to the interferon alpha and gamma pathways had significantly lower expression in GF fetal brain. These were partly overlapping and partly different from the DE interferon stimulated genes in the intestine, but included known ISGs, such as Rsad2, Ifi27, Ifitm3, and Bst2. Rsad2, also known as viperin, was the most significantly DE gene in brain. Its expression was also significantly lower in GF fetal intestine and placenta. Rsad2 protein expression is induced by interferon, and it has a major role in antiviral defense of the cell (Rivera-Serrano et al., 2020). It is an enzyme catalyzing the production of ddhCTP which is a direct inhibitor of virus replication by acting as a chain terminator of RNA-dependent RNA polymerases. It may also have other mechanisms of antiviral action (Zhang et al., 2007).

The importance of gut microbiota on the brain resistance for virus infection was recently shown by Yang et al. who showed that depletion of gut microbiota in mice exacerbated the neurological symptoms of EMCV infection concurrently with diminished innate immune responses and decreased expression of ISGs in brain (Yang et al., 2021).

Maternal microbiota appears to have a major impact on fetal neurodevelopment. Genes coding for multiple key neuronal and glial transcription factors and other regulators, structural proteins and synaptic signalling components were differentially expressed in GF and SPF fetuses, with almost all synaptic genes (by Gene Ontology annotations) downregulated in GF fetal brain. Modulation of fetal neurodevelopment by maternal microbiome, likely involving microbial metabolites, was recently reported (Vuong et al., 2020). However, the differentially expressed genes were almost completely different, primarily affecting axonogenesis and not enriched for any type of immunity genes (not shown). Vuong et al. studied younger (E14.5) fetuses, suggesting that the maternal microbiota modulates fetal neurodevelopment at several different stages. Lack of maternal microbiota affected the microglia in E18.5 mice, especially in male fetuses (Thion et al., 2018); differential expression of *Rtp4, Ifi27, Ifitm3* and *Bst2* were observed also here. Immunity and neurodevelopment may be linked by the cytokine CX3CL1 (fractalkine, neurotactin), which was significantly downregulated in our GF fetuses (Arnoux and Audinat, 2015). In brain, the CX3CL1 receptor (CX3CR1) is exclusively expressed in microglia. In CX3CR1 deficient mice, synaptic pruning and signalling and brain connectivity are deficient, leading to disruption in social interaction and behavior (Zhan et al., 2014).

### Placenta

In placenta, the maternal microbiota broadly modulated genes associated with growth and morphogenesis, especially blood vessel development, in addition to immunity genes. Several critical components of placental function (epidermal growth factor receptor, prolactin, and a trypsin inhibitor involved in pregnancy maintenance) were among the most significantly differentially regulated genes in the GF placenta, suggesting that the normal maternal microbiota contributes to the healthy pregnancy. Interestingly, *Endod1*, coding for an exosomal protein, was among the genes most significantly upregulated by maternal microbiota. Extracellular vesicles are thought to mediate immunological interactions in placenta, and it is tempting to speculate that their production could be modulated by the maternal microbiota (Kaminski et al., 2019). The lncRNA Gm7932 upregulated in the SPF placenta was recently implicated in virus defense response (Zhao et al., 2018).

### Insulin degrading enzyme (Ide)

*Ide* was by far the most downregulated gene in all GF tissues. To our knowledge, this effect of the maternal microbiota has not been previously reported in the fetus. The GF dams probably had lower levels of circulating insulin (Bäckhed et al., 2007). Maternal insulin is thought not to cross the placenta, but it does stimulate placental nutrient transport (Ruiz-Palacios et al., 2017). Thus, the differential *Ide* expression in the fetus may be an indirect consequence of higher nutrient availability, which likely affects fetal insulin levels. However, *Ide* expression was also strongly and positively associated with the levels of several microbially modulated metabolites (including 3-indolepropionic acid in the intestine and TMAO in brain), suggesting that it may be more directly modulated in the fetus by the maternal microbiota. Interestingly, several *Ceacam* genes were also downregulated in our GF intestine (some of which were also observed by de Aguero et al. or by Garcia et al.). Ceacam1 is involved in both insulin signaling, (Kuespert et al., 2006), tolerogenic immunological signaling (Kim et al., 2019) and mucosal colonization (Voges et al., 2010), and may therefore connect these processes (Foley et al., 2020).

### Associations of microbial metabolites with fetal gene expression

Multiple metabolites modulated by maternal microbiota were strongly associated with gene expression profiles in the fetal tissues and placenta. These included previously identified microbial metabolites and potential novel compounds of microbial origin (Pessa-Morikawa et al., 2022). Some of these could be at least tentatively annotated; further studies are needed to annotate others.

Several aryl sulfates likely produced by maternal microbiota were associated with immunity pathways in fetal intestine and brain: 4-hydroxybenzenesulfonic acid, indoxyl sulfate, pyrocatechol sulfate or isomer, and other phenylsulfates. In fetal intestine they were also associated with lipid and tRNA metabolism and regulation of growth, and in brain, with dopaminergic synapse genes.

Another microbial tryptophan derivative, 3-indolepropionic acid (IPA), as well as kynurenine and tryptophan itself, were also associated with immunity and translation in the fetus.

The trimethylated compounds trimethylamine-*N*-oxide (TMAO) and 5-aminovaleric acid betaine (5-AVAB) were associated with immunity, lipid metabolism and absorption in the fetal intestine. TMAO shared many of the most significantly associated genes with aryl hydrocarbons. TMAO was previously shown to promote fetal thalamocortical axonogenesis (Vuong et al., 2020), but in our fetal brain samples, we only observed strong association with the expression of insulin degrading enzyme (*Ide*) gene. This may be due to the different developmental stages examined.

Short-chain fatty acids (SCFA) and secondary bile acids are well-known mediators of host-microbe interactions (Li et al., 2022). SCFAs were not optimally detected by our metabolomics pipeline. However, the carnitine conjugate of butyrate was among the metabolites most significantly associated with immunity and lipid metabolism genes in the intestine.

A bile acid matching chenodeoxycholic acid was strongly associated with immunity, translation and intestinal absorption genes.

Some of the microbially modulated molecular features with strong associations with gene expression could not yet be annotated. A molecular feature with the probable formula C3H4O5 (possibly 2-hydroxypropanedioic acid, also known as tartronic acid or hydroxymalonate) showed the largest number of associated genes, enriched for immunity pathways. Multiple unannotated features were associated with neuron projection and myelin sheathing in fetal brain. Metabolites putatively annotated as N-linoleyltyrosine or its isomer, and peiminine or N-oleoylphenylalanine showed multiple strong associations with gene expression in the intestine.

Associations of microbially modulated metabolites with immunity related genes were mostly positive, and with translation related genes mostly negative. However, aryl sulfates, TMAO, betaines, bile acids and certain other metabolites showed mostly negative correlations with immunity genes, which was surprising considering that many of these compounds are generally considered proinflammatory. In GF fetuses they were associated with unexpected gene expression profiles: when gene expression was negatively correlated with metabolite signal in SPF fetuses, the GF fetuses showed *lower* expression than SPF fetuses for most of these genes; for positively correlating genes, the GF expression levels were *higher*. These genes were enriched for *negative* regulation of immunity (whereas genes with more linear correlation in GF fetuses were enriched for *positive* regulation). Our observations may suggest that these microbial metabolites downregulate anti-inflammatory genes; in GF fetuses, these pathways may be inactive in the absence of microbial proinflammatory signals.

Several aryl hydrocarbons were strongly associated with genes enriched for predicted binding sites of aryl hydrocarbon receptor (AhR) (Gao et al., 2018) or vitamin D receptor (VDR), and for known target genes of these and other xenobiotic-sensing nuclear receptors like pregnane X receptor (Venkatesh et al., 2014). However, some of the canonical AhR target genes (such as those coding the cytochrome P450 family 1 enzymes) were not significantly differentially expressed in GF and SPF fetuses. This suggests that the effects of these compounds in the fetus may be mediated by non-canonical AhR signalling or other xenobiotic signaling pathways.

### Consequences of fetal exposure to microbial metabolites

Early host-microbe interactions are essential for proper development of the immune system, metabolism and even central nervous system (Ganal-Vonarburg et al., 2020). These interactions begin already *in utero*, but the significance of the first encounters with microbes or their products are poorly understood. Fetal exposure maternal microbiota and bacterial AhR ligands is necessary for the differentiation of intestinal group 3 innate lymphoid cells and balanced intestinal immunity (Gomez de Aguero et al., 2016; Lu et al., 2021). Of the microbial aryl hydrocarbons identified in our fetal mice, indole-3-propionic acid is considered a neuroprotective antioxidant, which also improves the intestinal barrier (Vanholder et al., 2022). Indole-3-acetic acid may promote tissue repair and improve inflammation.

Our data indicates that the fetus is also exposed to several microbial metabolites which are often considered harmful. Indoxyl sulfate, kynurenine and indoleacetic acid are well-known uremic toxins in the context of kidney diseases (Vanholder et al., 2022), contributing to inflammation, cardiovascular diseases, and metabolic, hormonal and neuronal dysfunction. Another aryl sulfate (which we did not detect in our data), 4-ethylphenyl sulfate (4EPS), was recently reported to impair oligodendrocyte maturation and myelination in mice and to induce anxiety-like behavior (Needham et al., 2022). TMAO and its bacterially produced precursor TMA have been widely implicated in metabolic pathologies including atherosclerosis, metabolic disease, human type 2 diabetes and gestational diabetes (Krueger et al., 2021). Multiple biological mechanisms of action have been demonstrated in human and mouse models (Chen et al., 2019; Krueger et al., 2021), with the molecular roles of these compounds strongly dependent on the experimental setting and context.

Animals evolved under constant exposure to microbial metabolites, and the developing fetus is expected to tolerate or even require also these compounds. TMAO abrogated defects of axonogenesis observed in embryos of antibiotic-treated mouse dams (Vuong et al., 2020). Evaluating the physiological consequences of fetal microbial metabolite exposure and immune activation mediated by these compounds requires studies in postnatal animals. It is also context dependent: while germ-free animals are obviously abnormal in some aspects, other implications of the deficits in early host-microbe interactions are only realized in the face of environmental stressors. We are therefore studying the early host-microbe interactions in large production animals, in addition to laboratory mice living in strictly controlled conditions (Hamilton et al., 2020).

### Limitations of the study

This is an explorative study; *in vitro* and *in vivo* experiments with purified compounds identified here are necessary to show actual causal effects of microbial metabolites on gene expression and development. Meaningful statistical significances could not be computed for the observed associations between metabolites and gene expression. Over the whole dataset (including both SPF and GF animals), statistical significance is to be expected, as we examine associations of differentially expressed genes and differentially abundant metabolites. Within the SPF group, the observed associations were largely not statistically significant, due to the large number of metabolomics signals and genes, and the limited number of data points. However, we only took into account multiple very strong correlations (Spearman ρ>0.9) generating highly significant hits in over-representation analysis; we believe this to be a reliable approach to identify the metabolites which are most likely to be physiologically relevant.

Some of the differences in germ-free and control fetuses may be due to the abnormal physiology of the germ-free mouse dams, rather than direct effects of maternal microbial metabolites. However, this does not affect the reported associations of microbial metabolites and gene expression profiles, as these were examined in the SPF fetuses.

Gene expression profiling obviously does not capture all potential impacts, such as possible modulation of the antigen receptor repertoires of the adaptive immune system which requires macromolecular ligands.

## Conclusions

- Maternal microbiota has a major impact on the developing fetus. Immune system, neurophysiology, translation and energy metabolism are strongly affected already before birth.
- The impact is especially pronounced in the fetal intestine. Also in the developing brain, the impact of microbial metabolites appears substantial, although less profound, possibly due to the blood-brain barrier which limits the metabolite exposure of the neural tissue.
- These impacts are largely, although probably not exclusively, mediated by microbial metabolites, including still unidentified compounds.
- Several aryl sulfates are among the metabolites strongly associated with fetal gene expression.
- The germ-free fetus may suffer from depletion of queuine, the bacterial hypermodified nucleobase essential for eukaryotic tRNA stability and function.

## Acknowledgements

RNA sequencing was performed by the Sequencing unit of Institute for Molecular Medicine Finland FIMM Technology Centre, University of Helsinki. The sequencing unit is supported by Biocenter Finland. We thank Kirsi Lahti and Santeri Suokas for expert technical assistance.

## Ethical statement

The breeding and organ collection upon euthanasia are not considered experimental animal procedures (European Directive 2010/63/EU and Portuguese Law) and specific permission was not required.

## Author contributions

MN, AI and KH designed the study. AH performed the RNASeq data processing and differential gene expression analysis, TPM did the GSEA analyses, and MN did the ORA and association analyses. VK performed the metabolomics data processing and metabolite identification; OK and KH supervised the metabolomics analysis. All authors contributed to data analysis and interpretation. MN, TPM, AH and AI wrote the manuscript. All authors read, commented, and approved the final version of the manuscript.

## Data availability statement

The metabolomics dataset from (Pessa-Morikawa et al., 2022) can be found at EUDAT: https://doi.org/10.23728/b2share.4be0ea9f87b84a06be960d6a1c4b0b42. RNASeq data will be available upon publication of this study.

