## Supplementary Figures for "Impacts of maternal microbiota and microbial metabolites on fetal intestine, brain and placenta"

CC-BY 4.0

**Supplementary Figure 1. Heatmaps for the interferon alpha and interferon gamma response gene sets.**
a) Intestine Hallmark Interferon alpha Response, b) Intestine Hallmark Interferon gamma Response, c) Brain Hallmark Interferon alpha Response, d) Brain Hallmark Interferon gamma Response

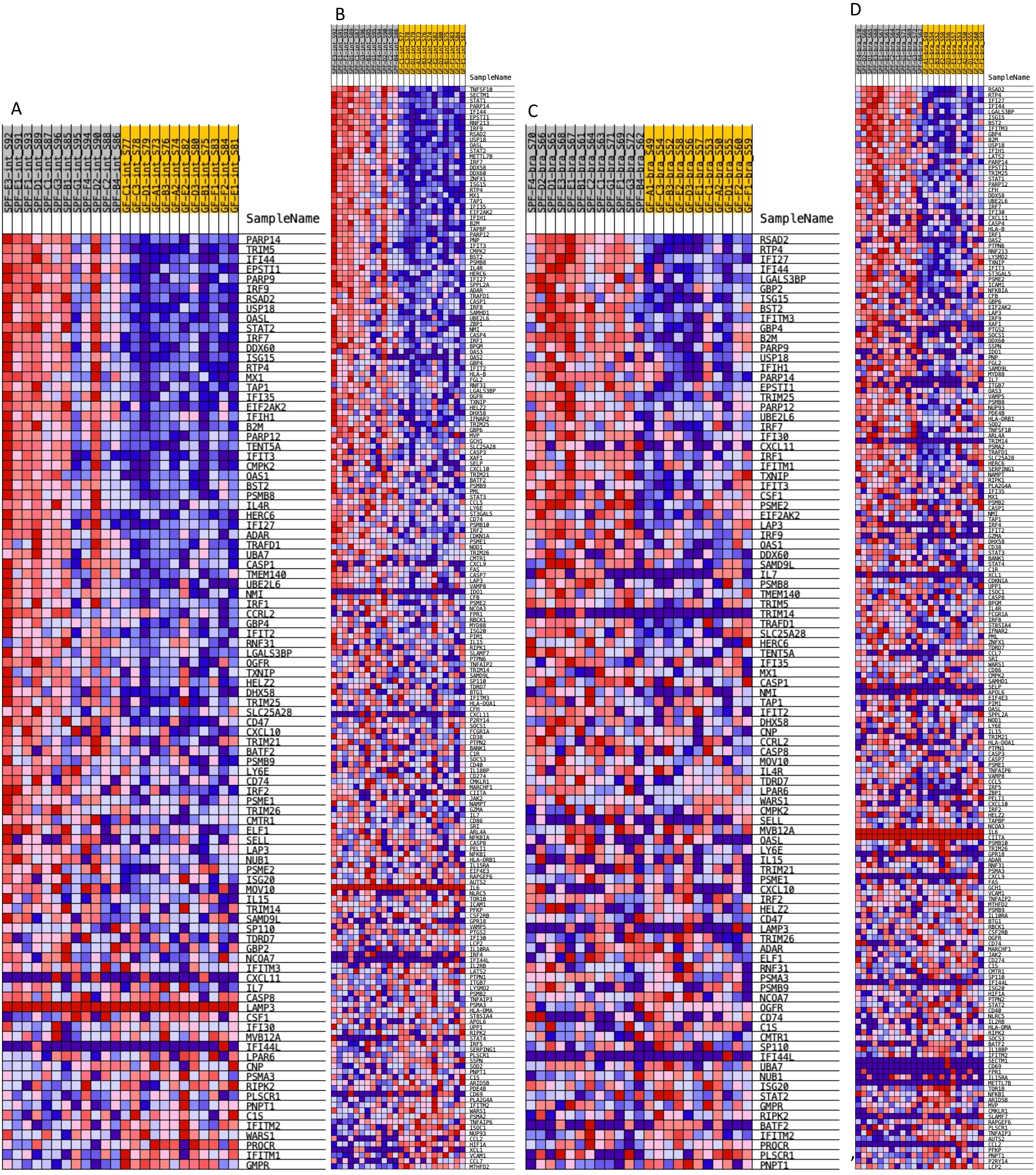

**Supplementary Figure 2. Over-representation analysis of genes which were significantly differentially expressed (q < 0.05) in GF versus SPF fetal murine intestine, brain and placenta. Top 100 enriched ontology terms are shown.**

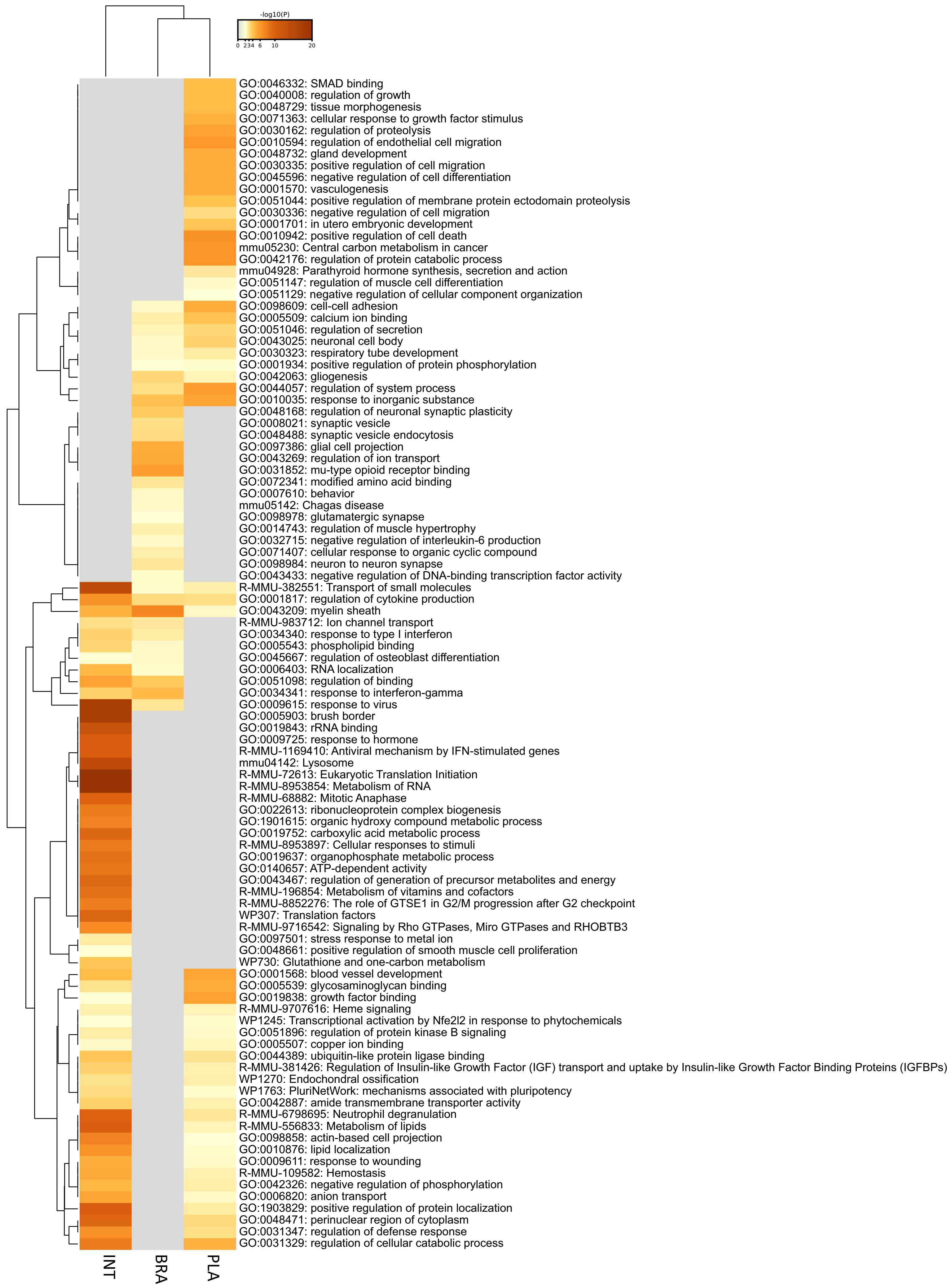

**Supplementary Figure 3. Over-representation analysis of genes strongly associated with metabolites in fetal intestine.** Highest-scoring molecular features missing from GF fetuses (left) and highest-scoring annotated metabolites more abundant in SPF fetuses (right). Top 100 enriched ontology terms are shown. Here, non-selective picking of ontology clusters was used.

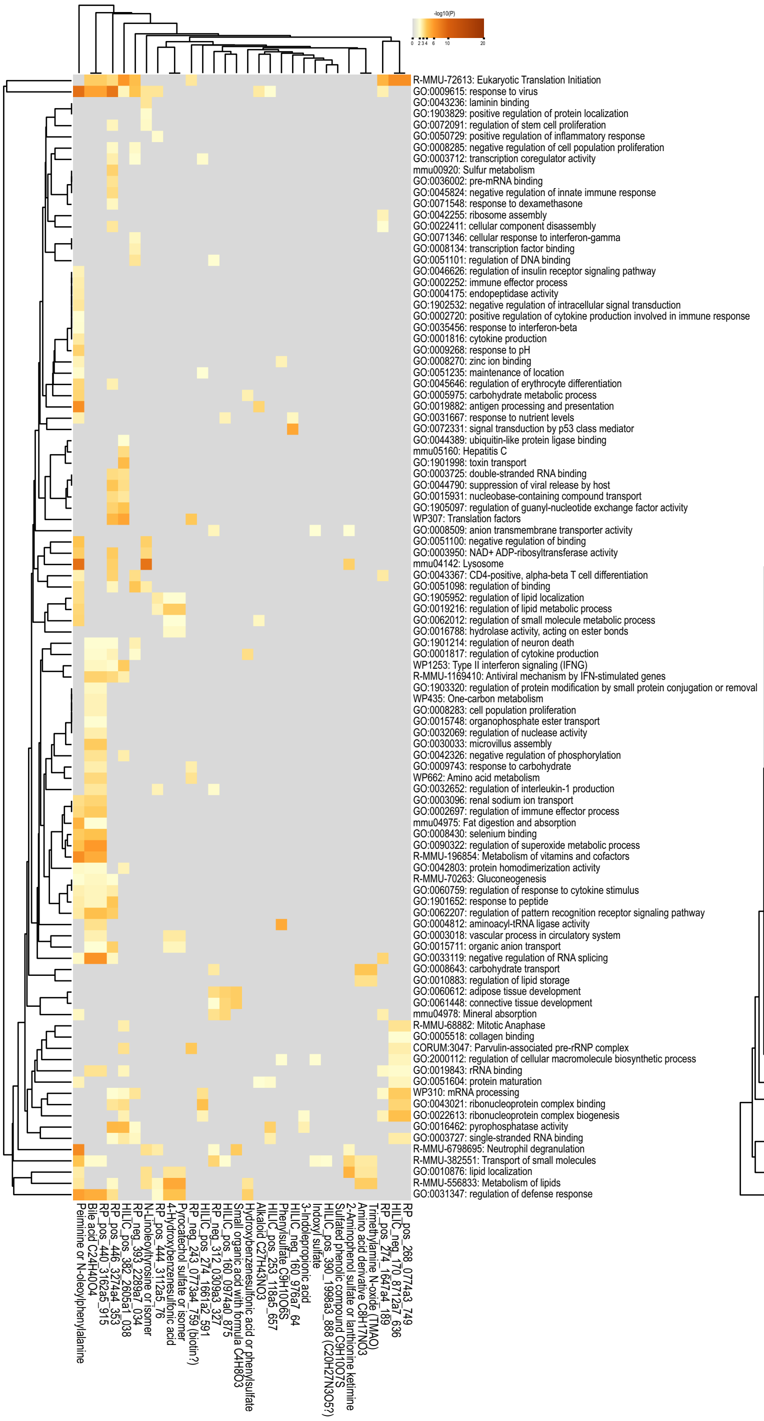


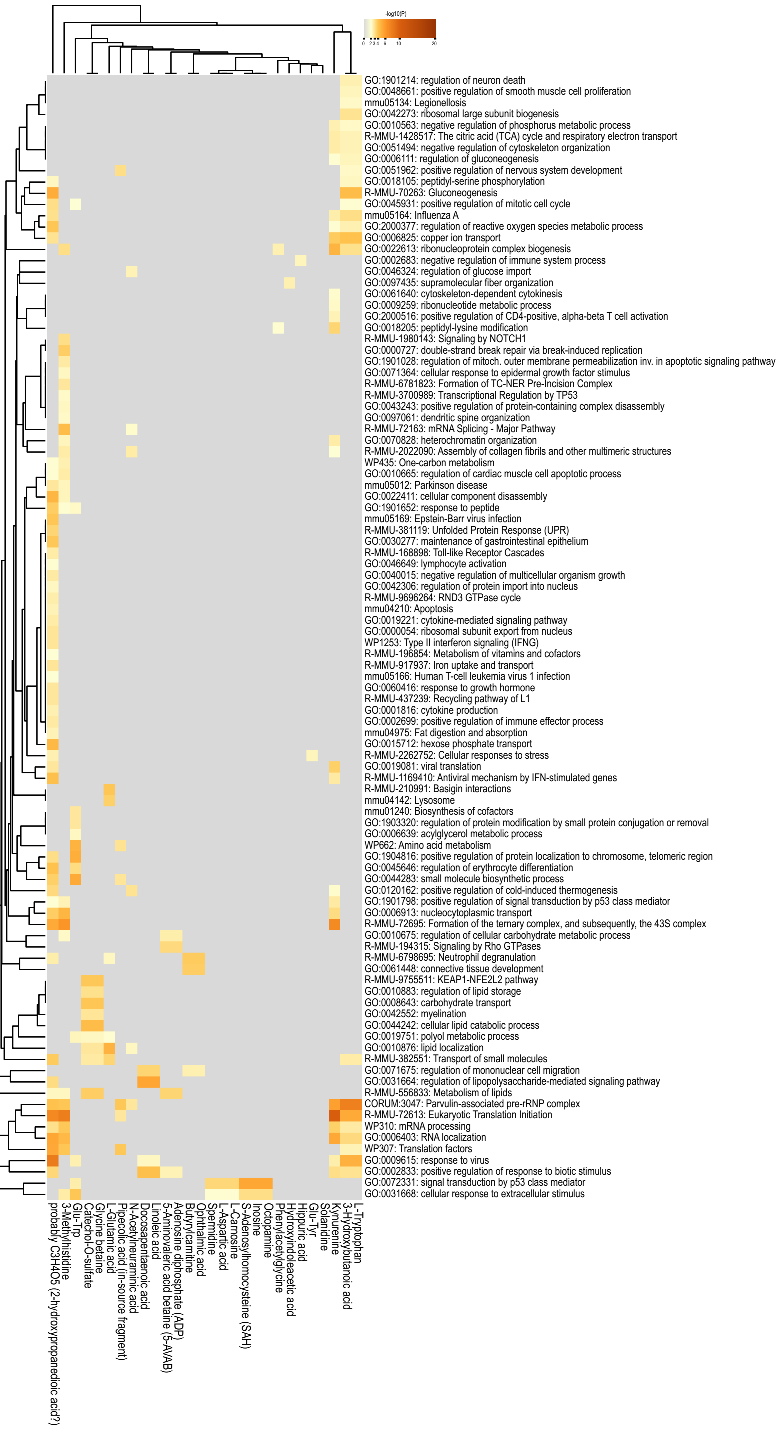

**Supplementary Figure 4. Over-representation analysis of genes strongly associated with metabolites in fetal brain.** Metabolites missing from GF fetuses are indicated by *. Top 100 enriched ontology terms are shown. Here, non-selective picking of ontology clusters was used.

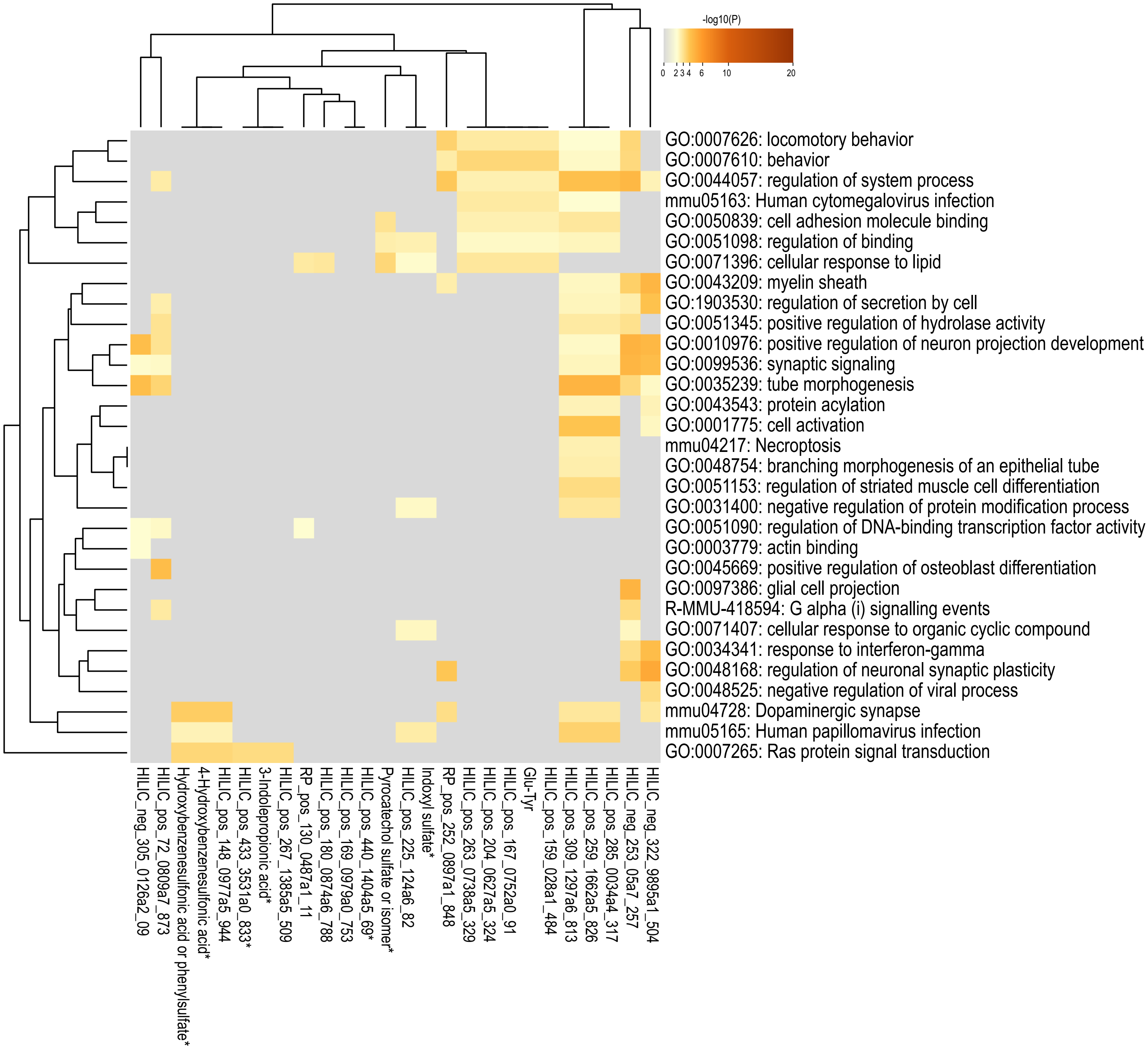

**Supplementary Figure 5. Over-representation analysis of genes strongly associated with metabolites in placenta.** Highest-scoring molecular features missing from GF fetuses (top) and highest-scoring annotated metabolites more abundant in SPF fetuses (bottom). Here, non-selective picking of ontology clusters was used.

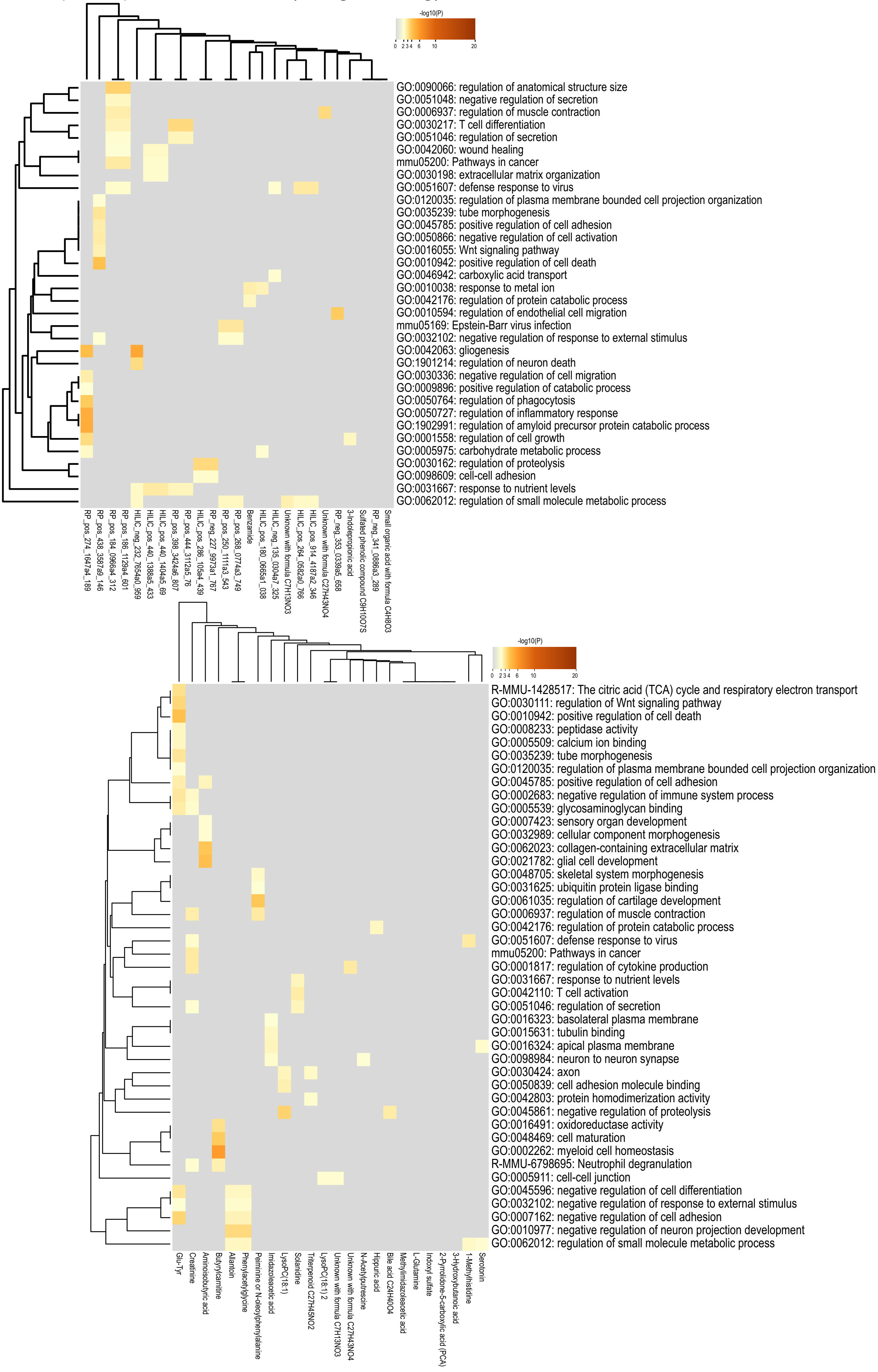
